# Sensory-motor integration in a nonspiking interneuron contributes to active sensor control in *Drosophila*

**DOI:** 10.64898/2026.04.16.718965

**Authors:** Olivia M. Nunn, Kaylee M. Odum, Ava C. Thorsen, Marie P. Suver

## Abstract

An animal’s nervous system enables it to detect and respond to stimuli to navigate its environment. To enhance sensory acquisition, animals can actively position sensors, altering how they extract information from the external world. However, active sensing, and movement in general, produces sensory feedback, requiring mechanisms for integrating predictive motor signals with externally generated sensations. Despite the importance of these mechanisms for guiding coordinated behavior, the cellular and circuit basis of motor control and sensory processing during active movements are not fully understood. Here, we investigate how mechanosensory information contributes to motor output in the *Drosophila* antenna, a sensor that can be actively positioned. We combine electrophysiology, quantitative behavior, optogenetics, and connectomics to characterize APN2, a nonspiking interneuron in *Drosophila* that encodes both mechanosensory stimuli and motor commands. We show that these neurons receive input from two classes of antennal mechanosensors, enabling responses to mechanical perturbations in both quiescence and flight. We then demonstrate how these neurons integrate higher-order motor commands with mechanosensory input to shape antennal movement. Together, this work reveals a previously uncharacterized sensory–motor circuit in the antennal mechanosensory system. These findings provide insight into how nervous systems integrate sensation with motor commands to guide the movement of active sensors, highlighting circuit mechanisms that may broadly support sensory acquisition during movement.

## Introduction

Animals often actively move their sensory organs to shape how they sample information from the environment. This strategy, known as active sensing, requires neural circuits that integrate sensory input with motor commands controlling sensor position. For example, humans use their fingertips to actively probe surfaces to determine the size, texture, and location of objects^1,2^. Similarly, rodents use their whiskers to actively sense their tactile environment^3–5^. This strategy extends to arthropods, which use their antennae – a pair of mobile, multi-modal sensory appendages – to extract mechanosensory information from the external world^6^. Despite the widespread importance of active sensing behaviors, the neural circuit basis for active sensor control is not fully understood.

Like other organisms, fruit flies actively position sensors to alter sensory acquisition. Their antennae are a bilateral pair of highly multimodal sensors actuated by a set of four muscles at their base^7,8^. As one of the primary mechanosensory appendages in the animal, the antennae house the largest mechanosensory organ in the animal, composed of ∼1000 primary mechanoreceptors called Johnston Organ Neurons (JONs)^9^. These bipolar sensory neurons convert mechanical forces into electrical signals, with different subsets of JONs encoding antennal movements produced by sound, wingbeats, and airflow^10–14^. All JONs primarily project to the antennal mechanosensory and motor center (AMMC) in the central brain, where they synapse onto AMMC projection neurons that encode stimulus features including frequency, amplitude, and phase^13^.

Antennal sensory circuits play a key role in many behaviors in flies. For instance, during flight, mechanosensory information detected by the antennae contributes to steering^11^, groundspeed regulation^15^, and odor tracking^16–18^. In addition to passively detecting airflow, recent work shows that active antennal movements can contribute to these behaviors by tuning wind encoding^8^. Moving the antennae will likely influence sensation by sensory neurons housed in the antennae, so how do antennal sensory-motor circuits coordinate ongoing sensory input with motor commands to control active movements?

One candidate mechanism for sensory-motor integration is graded signaling by nonspiking interneurons, which often serve as an intermediate between sensory afferents and motor neurons in arthropods^19–21^. Unlike all-or-nothing spiking activity, graded changes in membrane potential preserves fine stimulus features during transmission from sensory afferents to motor circuits^22^. Graded changes in membrane potential control rhythmic motor output^23,24^, posture maintenance^25,26^, and reflex modulation^27,28^. More recently, work in the *Drosophila* leg motor circuits identified a nonspiking interneuron that mediates slow postural leg movements in response to limb perturbations detected by the femoral chordotonal organ^29^ – a functionally analogous structure to the Johnston’s Organ in the antennae^30^. Whether antennal mechanosensory pathways similarly employ nonspiking interneurons to translate stimulus features like frequency, amplitude, and phase from the JONs to antennal motor circuits controlling active behaviors has yet to be determined.

In this study, we study the role of AMMC projection neuron 2 (APN2), a nonspiking interneuron in the *Drosophila* antennal mechanosensory circuit^31,32^, and find that it contributes to active antennal positioning via integration of sensory and higher-order motor signals. Using whole-cell patch clamp electrophysiology in tethered behaving flies, we show that APN2 activity is modulated during flight by inhibitory input, suggesting a circuit mechanism that could adjust antennal posture during behavior. Despite this flight-dependent modulation, APN2 remains responsive to external airflow through two distinct antennal mechanosensory pathways. We further demonstrate that APN2 activity is correlated with active antennal movements and, using optogenetics, show that it contributes to antennal motor output. Finally, we outline the broader sensory–motor circuit in which APN2 operates, using connectomics data to link antennal mechanosensory input to downstream motor neurons. Together, these findings define the role of APN2 in antennal sensory–motor integration and provide a framework for understanding how neural circuits use sensory information to guide active sensor movements.

## Results

### A second-order mechanosensory neuron is hyperpolarized during flight

Wing-generated airflow during flight produces ipsilateral antennal oscillations, activating JONs^12^. To examine whether increased JON activity during flight influences downstream neural responses, we recorded intracellular activity of APN2, a class of neurons in the central brain that are postsynaptic to JON subclasses C and E and respond to externally generated mechanical stimuli^31,32^ (Figure 1A). We performed whole cell patch clamp recordings in intact flies while simultaneously recording ipsilateral antennal position and wingbeat activity (Figure 1B-D). In these recordings, APN2 exhibited a sustained decrease in membrane potential during flight relative to quiescence (Figure 1D-E). During quiescence, APN2 baseline membrane potential was −34.1 ± 2.0 mV, whereas during flight it dropped to −36.4 ± 1.7 mV (Figure 1E; paired t test, p = 0.037), 2.3 mV more hyperpolarized than during quiescence. To test whether this sustained decrease in activity depended on input from JONs, we mechanically blocked JON activity by applying glue to the A2/A3 joint on both antennae^33^. Blocking JON activity did not abolish the hyperpolarization during flight, but it did greatly reduce its magnitude (Figure 1E). On average, APN2 was slightly but significantly more hyperpolarized during flight (−39.4 ± 0.9 mV) than during quiescence (−38.1 ± 1.0 mV; Wilcoxon signed-rank test, p = 0.014). We observed no significant differences between flies with free and JON blocked antennae during quiescence (independent t test, p = 0.085) or in flight (Mann-Whitney U, p = 0.193).

**Figure 1.**
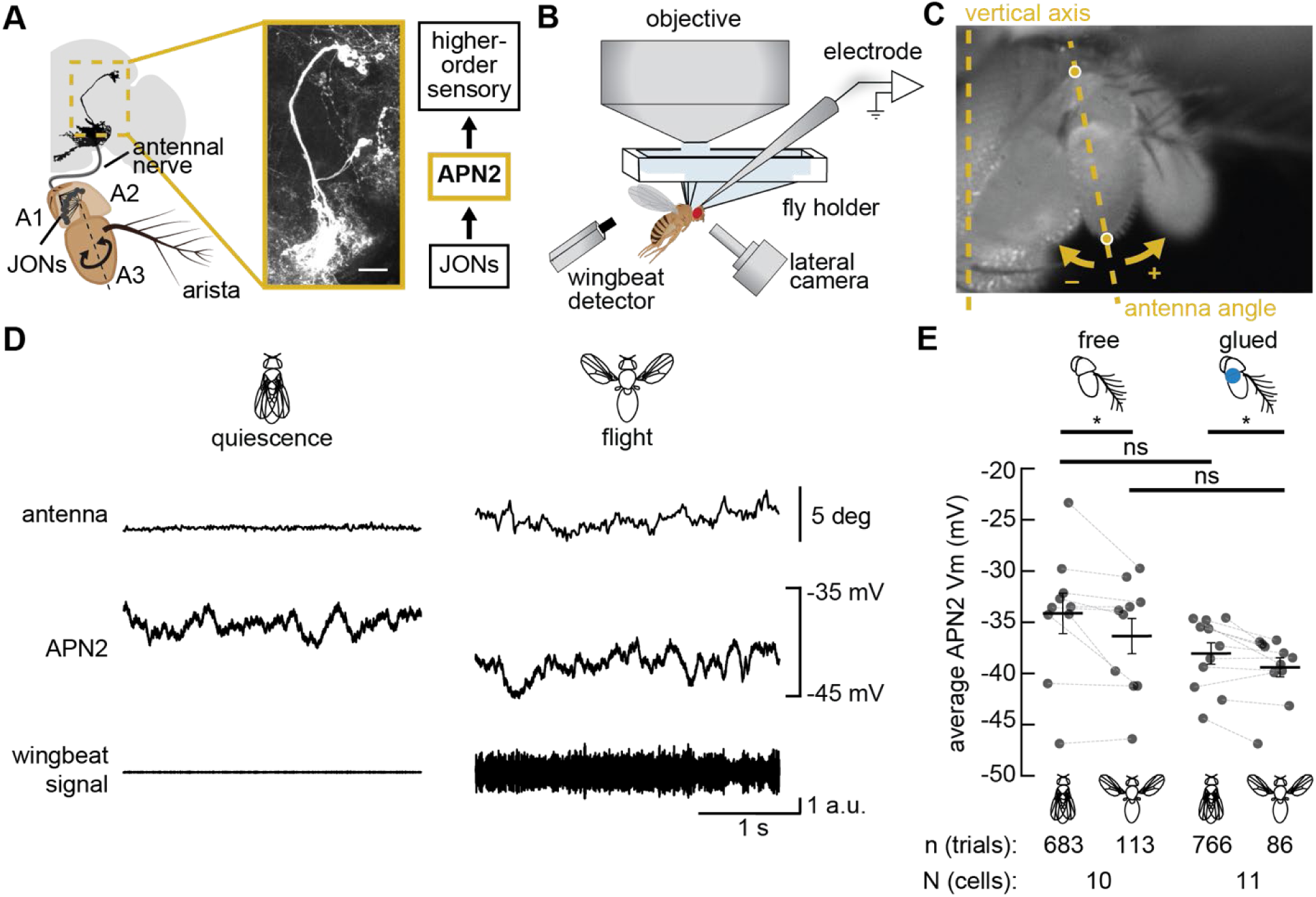
Flight induces sustained hyperpolarization in APN2, with and without JON input. A) Schematic depicting JONs projecting through the antennal nerve and synapsing onto APN2 cells (gold dashed box) in the ipsilateral hemisphere of the central brain. Inset shows expression pattern of the genetic driver line labeling APN2 (*24C06-GAL4)*. Scale bar is 20 µm. B) During whole cell patch clamp recordings, antennal movements were recorded by a lateral camera and flight activity was monitored using an optical wingbeat detector. C) Example video frame with 2 tracked points and the relative direction of antenna angle deflections. Deflections down towards the head are represented as negative values, and deflections up and away from the head as positive values. D) Example single-trial traces showing antennal and APN2 activity during quiescence (left) and flight (right). Wing movement was detected using an infrared light sensor. E) Each dot is the average response of one APN2 cell. Black bars indicate the average across APN2 cells for each condition, with respective SEM error bars. Grey dashed lines pair measurements from the same cell during flight and quiescence. Across flies, the average APN2 membrane potential is reduced during bouts of flight compared to quiescence, both when the antennae are free (paired t-test; p = 0.037) and when the antennae are glued (Wilcoxon Signed-Rank Test; p = 0.014).

### Mechanosensory input from JONs contributes to antennal posture during mechanical perturbations in quiescence and flight

In addition to airflow generated by the wings during flight, externally-generated mechanical stimuli – like wind – can increase JON activity^10,14^. To test whether increased JON activity affects antennal position, we delivered short air puffs (180-400 ms long) to the fly during both quiescence and flight while recording APN2 intracellular responses (Figure 2A). We then compared antennal responses to these short pulses of airflow in flies with intact versus blocked JON input^33^ (Figure 2B-C). When JON input was blocked, we observed an increase in the magnitude of the airflow-induced antennal deflections (Figure 2C). To quantify this, we plotted the correlation between antennal position and APN2 activity during an air puff (Figure S2), comparing quiescence and flight both with (Figure S2A) and without JON input (Figure S2B). The distribution of airflow-induced antennal positions was broader during flight than during quiescence, both when the antennae were free (pairwise Levene’s test; p < 0.001) and when JONs were blocked (p = 0.012). The largest increase in the distribution of antennal position occurred when we blocked the JONs – both in quiescent trials (p < 0.001), and flight trials (p<0.001). Taken together, antennal position exhibited significant variability in response to a brief air pulse across flying and quiescent flies, both when the antennae were free and when JONs were blocked (Figure S2C; Levene’s test; p < 0.001).

**Figure 2.**
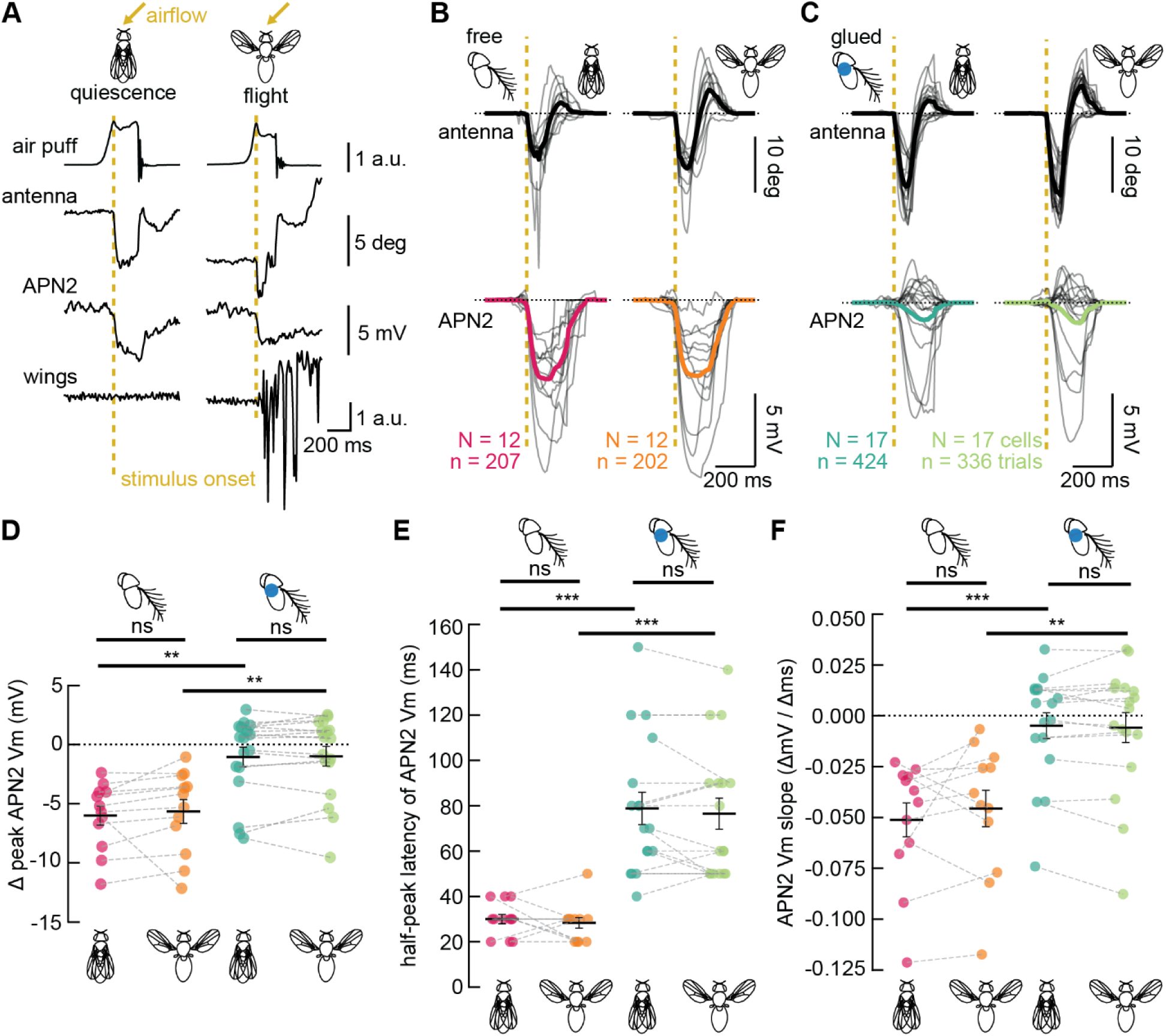
Mechanosensory responses in APN2 decrease, but are not fully eliminated, when JONs activity is blocked. A) Example single-trial traces showing airflow stimulus and the corresponding activity of the ipsilateral antenna, APN2 membrane potential, and wingbeat signal during quiescence (left) and flight (right). The gold arrow indicates the 45° angle of contralateral airflow delivered during each trial. The gold dashed line indicates when the air puff reaches the fly. B) Average responses to air puffs in quiescent and flying flies with free antennae. Each line is the average of all trials recorded in one APN2 cell. The thicker line represents the average across all cells. The horizontal dashed line represents the baseline antennal position (top) or the baseline membrane potential (bottom), and the gold vertical dashed line marks stimulus onset. C) Similar to (B), but when JONs are mechanically blocked. D, E, F) Each dot is the average of one APN2 cell. Black bars indicate averages across cells for each condition, with respective SEM error bars. Grey dashed lines pair measurements from the same cell during flight and quiescence. D) Blocking JONs reduced APN2 response amplitude of the airflow response in both quiescent and flying flies (Mann-Whitney U; p = 0.001 and p = 0.003, respectively). E) Blocking JONs increased APN2 peak latency in both quiescent and flying flies (Mann-Whitney U; p < 0.001 for both). F) Blocking JONs also decreased the slope of APN2 responses in both quiescent and flying flies, regardless of whether responses were hyperpolarizing or depolarizing (Mann-Whitney U; p < 0.001 and p = 0.002, respectively).

### APN2 receives input from two different antennal mechanosensors

Based on previous work^31,32^, we predicted that JONs would provide the majority, if not all, of the mechanosensory input to APN2. To test this, we compared APN2 activity in response to short pulses of airflow with intact and blocked JON input^33^, both in quiescence and in flight. When JON input is intact, APN2 membrane potential reliably hyperpolarized in response to airflow-induced antennal deflections (Figure 2B), consistent with previous work^31^. Surprisingly, when we blocked JON input, many APN2 cells (9/17 neurons) exhibited a depolarizing response to airflow (Figure 2C) - indicating another source of mechanosensory input to these cells. To characterize this response, we first measured the magnitude of the stimulus-evoked change in membrane potential by identifying APN2’s peak response (Figure 2D). Relative to the antennae-free condition, the amplitude of APN2’s airflow response was significantly reduced when JONs were blocked in both quiescent (Figure 2D; Mann-Whitney U; p = 0.001) and flying flies (p = 0.003), despite the absolute antennal deflection being considerably larger when the JONs were blocked (Figure 2C). Gluing the antennae decreased the peak APN2 response (Figure 2D) and delayed the time until peak response (Figure 2E). In flies with blocked JON input, the half-peak latency of the stimulus-evoked response was significantly prolonged across quiescent trials (Figure 2E; Mann-Whitney U; p < 0.001) and those with flight (p < 0.001). In both quiescent and flying flies, the response rate in APN2 from stimulus onset to peak (Figure 2F) became significantly slower when the JONs were blocked (Mann-Whitney U; p < 0.001 and p = 0.002, for quiescence and flight, respectively). Notably, all stimulus-evoked responses in APN2 were similar between quiescent and flight trials, regardless of JON input (Figure 2D-F), with little change in distribution (pairwise Levene’s test; free: p = 0.329, glued: p =0.961; Figure S2A-B). These results demonstrate that mechanical perturbations evoke consistent responses in APN2 during quiescence and flight, even without JON input.

### APN2 response magnitude increases during self-generated antennal movements

*Drosophila* produce self-generated antennal movements, likely altering sensation^8^. To explore how spontaneous antennal movements might influence APN2 activity, we next compared changes in membrane potential during spontaneous antennal movements to baseline activity recorded when the antennae were still (Figure 3). To further separate mechanosensory from motor-driven changes in APN2 activity, we compared APN2 responses in flies with intact (Figure 3A) and blocked JONs (Figure 3B). Because the antennae can move independently^8^, we tracked the position of both the ipsilateral and contralateral antennae while recording APN2 membrane potential. Previous work showed that APN2 receives ipsilateral antennal mechanosensory input^31^, so we hypothesized that it would not respond to contralateral antennal deflections. Unexpectedly, APN2 responded to both ipsilateral and contralateral antennal movements (Figure 3 A-B). To compare APN2 responses during inactive and active antennal states, we pooled ipsilateral and contralateral movements together (Figure 3C). In animals with free antennae, we measured spontaneous antennal movements in only 6 of the 12 recorded cells. Across these 6 cells, we detected a total of 14 active bouts. However, when we blocked JONs, we observed more spontaneous active antennal movements (in 12/17 cells). Across these 12 recordings with spontaneous movements, we detected a total of 59 active bouts. Together, we observed spontaneous active bouts in 50% of the recordings when the JONs were intact, compared to 70.6% when the JONs were blocked. Furthermore, the average number of active bouts per recording increased from 2.33 when the antennae were free to 4.92 when the antennae were glued – more than doubling the number of these spontaneous, self-generated movements. Not only did blocking JONs increase active antennal movements, but it also changed the activity of APN2. Active antennal movements evoked both depolarizing and hyperpolarizing responses in APN2, so we quantified its activity by calculating the absolute area under the continuous membrane potential response. When the antennae were free, the absolute area under the curve increased during spontaneous antennal movements compared to bouts of stillness (Figure 3C, Wilcoxon signed-rank test, p = 0.007). A similar trend held when the JONs were blocked, although the difference was not statistically significant (p = 0.088). Further, we observed significantly more activity when we compared activity in APN2 during active movements when the JONs were intact versus when they were blocked (Figure 3C, Mann-Whitney U, p = 0.010). Together, these data show that APN2 activity is modulated by spontaneous antennal movements and is shaped by mechanosensory input from the JONs.

**Figure 3.**
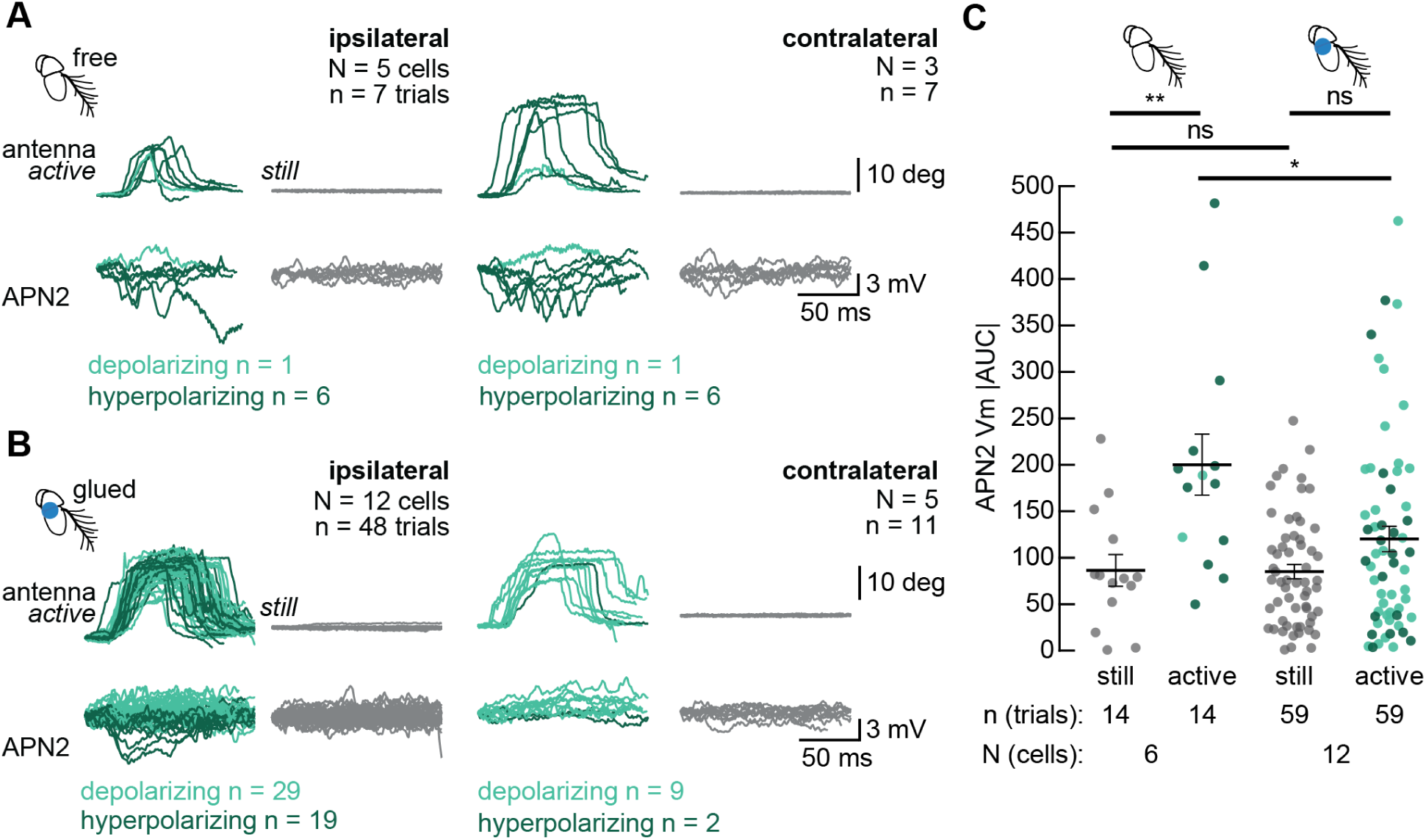
APN2 response magnitude increases during spontaneous active antennal movements. A) Single-trial traces showing spontaneous ipsilateral (left) and contralateral (right) antennal movements relative to the brain hemisphere in which we recorded APN2 activity in non-flying flies (APN2 membrane potential plotted below). Antennal movements corresponding to APN2 depolarization are plotted in light green, and those corresponding to hyperpolarizing responses are in dark green. For comparison, an equal number of trials from the same cells are plotted in grey to represent baseline activity when the antenna is still. B) Similar to (A), but when the JONs are mechanically blocked. C) Absolute APN2 membrane potential for every active antennal movement for both ipsilateral and contralateral responses. Black bars indicate averages across trials for each condition, with respective SEM error bars. The absolute area under the curve, or |AUC|, during active antennal movement is higher than during still periods, in antennae free and glued conditions (Wilcoxon Signed-Rank; free p = 0.007; glued p = 0.088). The absolute area under the curve during active movement is also higher when the antennae are free than when they are glued (Mann-Whitney U; p = 0.010).

### APN2 activation is sufficient to drive antennal movements

To determine if APN2 has a functional role in controlling antennal movements, we measured the change in antennal position in tethered flies when APN2 was optogenetically inactivated or activated (Figure 4A-C). We looked at the effect of optogenetic inactivation (Figure 4D-E) and activation (Figure 4F-G) of APN2 on antennal position in both quiescent and flying flies. Exposing control flies to inactivating (505 nm) light resulted in small ventral antennal deflections both during quiescence and flight (Figure 4D, top row). However, we did not observe this response when we inactivated APN2 with the same light stimulus (Figure 4D, bottom row). Instead, we observed small dorsal antennal movements (< 2 deg, on average) when we inactivated APN2, both during quiescence and flight (Figure 4E; t-test; p < 0.001 and p = 0.010, respectively), suggesting a role for APN2 activity in driving antennal movements. The change in antennal position in response to the light stimulus did not differ significantly between quiescent and flying animals, either in controls or during optogenetic inactivation of APN2 (Figure 4E).

**Figure 4.**
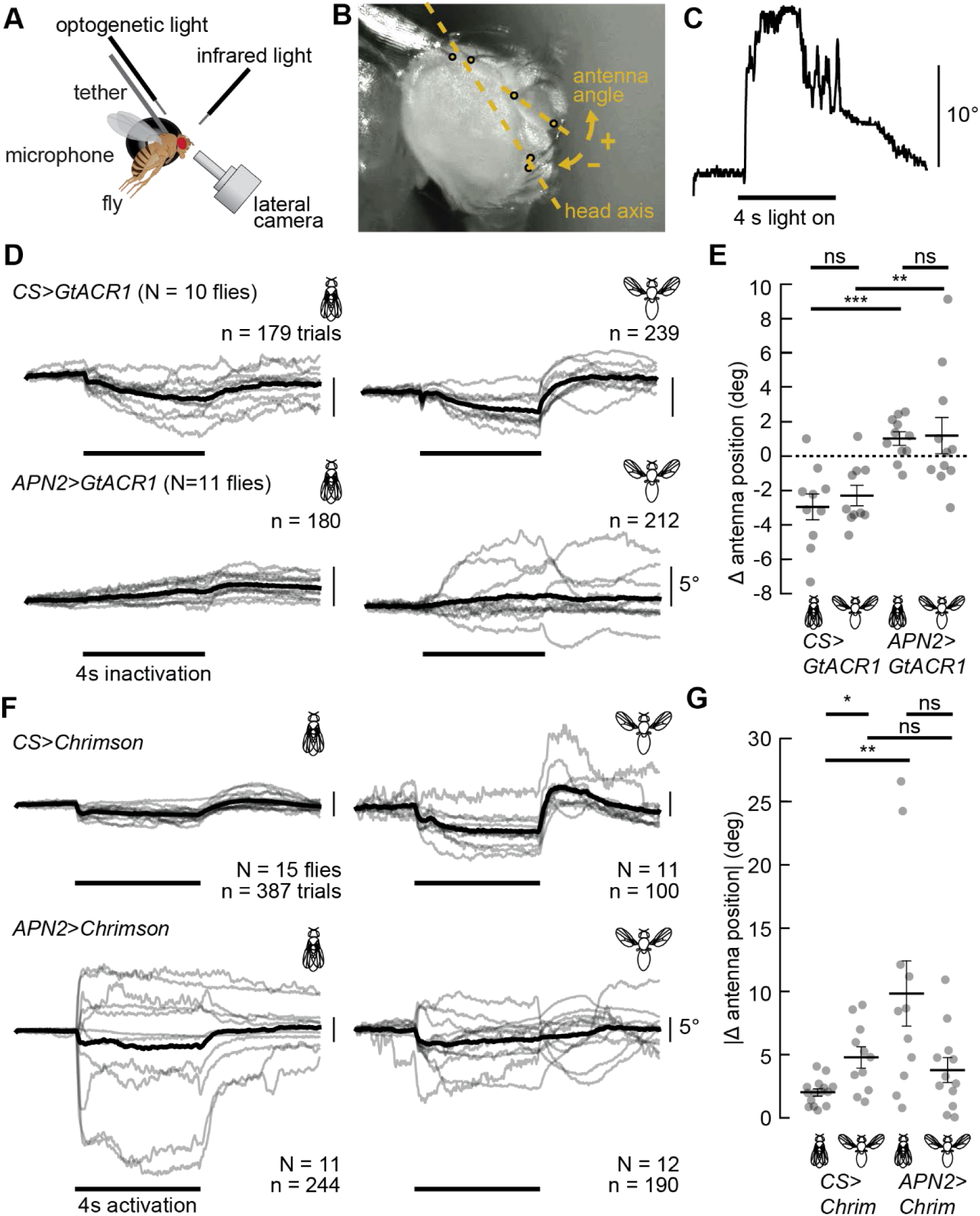
Optogenetic activation of APN2 elicits bidirectional antennal movements, whereas inactivation produces small upward movements. A) The optogenetic stimulus was presented to each fly using a fiber optic aimed at the fly’s head, antennal movements were recorded by a lateral camera, and flight activity was monitored using a microphone. B) Example video frame with 6 tracked points and the relative direction of antenna angle deflections. Deflections towards the head are represented as negative values, and deflections away from the head as positive values. C) Example single-trial trace showing antennal activity during APN2 optogenetic activation. D) Comparison of average antennal responses to 505 nm light in control (*Canton-S>UAS-GtACR1*) and APN2 (*24C06-GAL4>UAS-GtACR1*) flies during quiescence (left) and flight (right). Responses of control flies are plotted above and the effect of optogenetic inactivation of APN2 are plotted below. Each gray trace is the average of all trials in each fly, and the black traces represent the average across all flies. E) The change in antennal position during optogenetic stimulus was quantified by subtracting the average position before the stimulus from the average position during the last 2 s of stimulus. Each gray circle is one fly. Black bars represent the average across all flies and are plotted with their corresponding SEM error bars. In comparison to the control, inactivating APN2 decreased antennal movements in both quiescent (t-test; p < 0.001) and flying (t-test; p = 0.010) flies. F) Similar to (D) but using 590 nm light for controls (*Canton-S>UAS-CsChrimson*) and for the optogenetic activation of APN2 (*24C06-GAL4>UAS-CsChrimson)*. G) Similar to (E) but comparing the absolute change in antennal position observed in (D). Absolute antennal position during flight significantly increased during activation in control flies, but not APN2-activated flies (t-test; p = 0.037 and p = 0.053, respectively). In comparison to the control, activating APN2 increased antennal movements in quiescent flies (t-test; p = 0.004).

Like control flies in the inactivation experiments, we observed small ventral antennal deflections in control flies in response to the activating (590 nm) light stimulus (Figure 4F, top row). However, when we optogenetically activated APN2, we observed changes in antennal position larger than 25 degrees during quiescence and 10 degrees during flight (Figure 4F, bottom row). Because activating APN2 produces two roughly opposite directions of antennal movement (dorsal and ventral), we compared the absolute value of the change in antennal position resulting from the optogenetic stimulus (Figure 4G and S3). Compared to the control, we observed a significant increase in absolute antennal position during optogenetic activation of APN2 during quiescence (Mann-Whitney U; p = 0.004; Figure 4G), showing that activating APN2 is sufficient to drive antennal movements. We did not observe a significant change in the absolute antennal position during optogenetic activation of APN2 in flying flies compared to control flies (p = 0.601; Figure 4G).

### APN2 receives input from multiple antennal mechanosensors and is presynaptic to antennal motor neurons

We used the full adult female brain (FAFB) connectome^34,35^ to identify pre- and post-synaptic parters for APN2 (Figure 5A). Consistent with previous literature^31,32^, we identified strong connections (over 3,000 synapses) between JONs and APN2 within each hemisphere^34,36–38^. Both JONs and APN2 are likely cholinergic^34,37–39^ (Figure 5Ai), so we treated these connections as excitatory in our analysis. To focus on rapid sensory feedback to APN2, we limited our analysis to pathways containing a single interneuron between JONs and APN2. Of these sensory interneurons, 44.3% were inhibitory and 55.7% were excitatory (Figure 5Bii and Table S1). To identify candidates contributing to the JON-independent inhibition of APN2 we observed during flight, we analyzed all the additional inhibitory inputs into APN2 (Figure 5C). Independent of sensory input, we identified 4 inhibitory neurons that make 154 synapses onto APN2 (“flight-specific inhibition”, Figure 5Ai, Table S1 and S2), spanning both central and ascending super classes (Figure 5C).

**Figure 5.**
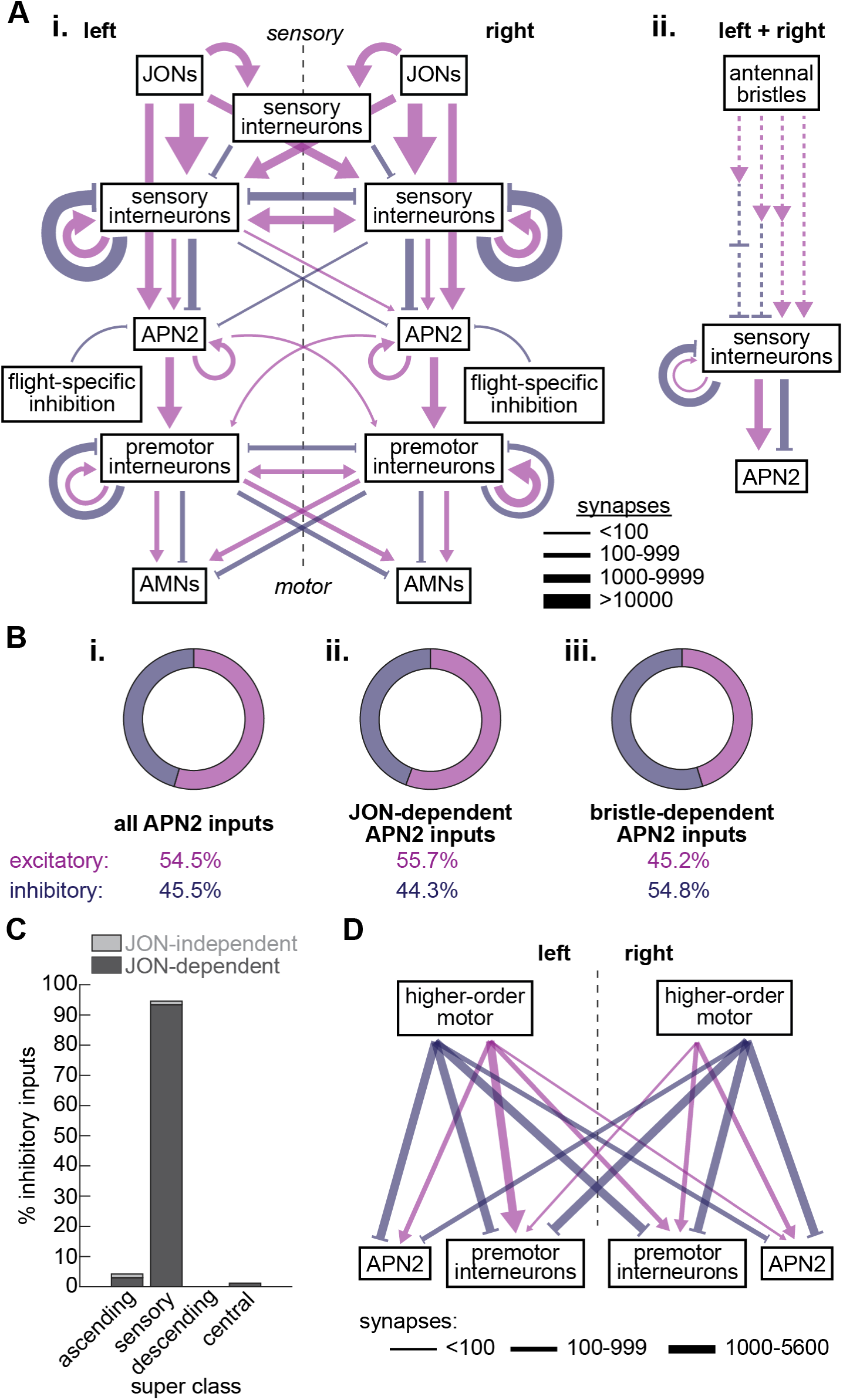
APN2 antennal sensory-motor circuit connectivity. A) A connectomics data set reveals direct and indirect sensory inputs to APN2 through (i) JONs and (ii) antennal bristles. The shortest pathways from APN2 to antennal motor neurons (AMNs) are via a single premotor interneuron. Excitatory connections are shown in pink, and inhibitory connections in purple. Solid lines indicate synaptic connections and are weighted on a log scale, and dashed arrows indicate synaptic connections that are not weighted due to the large number of possible intermediate pathways. B) Proportion of APN2 inputs that are excitatory versus inhibitory across (i) all inputs, (ii) JON-dependent inputs, and (iii) bristle-dependent inputs. C) Distribution of inhibitory inputs to APN2 by super class, categorized by whether the presynaptic neuron receives JON input (light gray), or do not (dark gray). D) Higher-order motor neurons project both ipsilaterally and contralaterally, providing input to APN2 and antennal premotor neurons.

We also investigated connections between bristles – another class of mechanosensors housed on the antennae^40^ – and APN2. We identified pathways consisting of one to three interneurons, totaling ∼2,500 additional mechanosensory synapses onto APN2 within each hemisphere (Figure 5Aii). Of the sensory interneurons within the bristle pathway that are directly presynaptic to APN2, 54.8% are excitatory, and the other 45.2% are inhibitory^34,37,38^ (Figure 5Biii and Table S1).

To understand how APN2 might provide feedback to the antennal motor system, we mapped pathways between APN2 and the antennal motor neurons labeled in the connectome (Figure 5Ai). We found pathways as short as one interneuron between APN2 and antennal motor neurons (Figure 5Ai). These premotor interneurons project to four of the ten antennal motor neurons via 4,962 synapses, with 45% of connections targeting contralateral motor neurons (Figure 5Ai, Table S1, and Table S2). Furthermore, we identified 210 neurons that cross the midline and project to both APN2 and premotor neurons (Figure 5D and Table S1), identifying candidate sources for higher-order motor signals coordinating between both antennae (Table S2).

## Discussion

To understand how sensory input influences motor commands controlling active sensor movements, we investigated APN2, a nonspiking class of interneurons that contribute to the encoding of mechanosensory stimuli^31^. Using whole-cell patch clamp electrophysiology, we show that APN2 exhibits a flight-dependent hyperpolarization (Figure 1) and encodes mechanical perturbations during both flight and quiescence using JONs and antennal bristles (Figure 2). APN2 also exhibits activity suggesting a premotor role: it is active during spontaneous antennal movements (Figure 3), and optogenetic activation drives antennal movements (Figure 4). Together with the sensory-motor circuit revealed through connectomic analyses (Figure 5), our findings suggest that APN2 integrates antennal mechanosensory input with motor control to support active sensing.

### Inhibitory control of APN2 during flight

Previous studies showed that JON activity increases in response to both external and self-generated mechanical stimuli^9,10,12,14^. Our work extends these findings by revealing how this activity shapes subthreshold neural activity in APN2 (Figures 1-2), which lies downstream of JONs tuned to tonic (or ‘push-pull’) stimuli^31,32^. Consistent with this connectivity, APN2 encodes tonic antennal deflections during quiescence^31^. However, during flight, JON subgroups positioned upstream of APN2 respond with an increase in activity during phasic oscillations of distal antennal segments^12^. Thus, we investigated APN2 cellular responses during flight, where we predicted that excitatory input from JONs activated by the vibrating wings would depolarize APN2. Unexpectedly, we observed sustained hyperpolarization in APN2 during flight (Figure 1E), suggesting additional flight-specific inhibitory input.

To determine whether the tonic decrease in APN2 membrane potential depends on increased JON activity, we mechanically blocked JON input and recorded from APN2 during quiescence and flight. Surprisingly, APN2 still exhibited a significant, though subtly reduced, hyperpolarization during flight when JON input was blocked (−1.4 mV; Figure 1E). This reduction was not statistically different than the hyperpolarization we measured when the JONs were intact (Figure 1E), indicating that although JONs contribute to the decrease in APN2 activity observed during flight, additional JON-independent inhibitory inputs likely drive this flight-specific hyperpolarization. Direct excitatory input from the JONs outnumber synaptic input from inhibitory sensory interneurons between JONs and APN2 (Figure 5Ai), so the inhibitory response during flight is unexpected given previous observations of a flight-dependent increase in JON activity^12^. However, 45.5% of APN2’s presynaptic partners in the connectome are likely inhibitory (expressing GABA or glutamate; Figure 5Bi). When the JONs are intact, the hyperpolarization of APN2 (Figure 2) is likely driven by the substantial recurrent inhibitory connections within the sensory interneuron population downstream of JONs – totaling over 10,000 synapses on either side of the brain (Figure 5Ai). When the JONs are blocked, the ascending inhibitory neurons that synapse directly onto APN2 (Table S2) likely contribute to the flight-specific hyperpolarization.

A sustained inhibition during flight could contribute to the control of antennal posture, supported by the upstream location of APN2 relative to antennal motor neurons (Figure 5A). During flight, the antennae are held in a more dorsomedial position than during quiescence, requiring mechanisms to maintain this posture during sustained activity. Previous work hypothesized that continuous neurotransmitter release from nonspiking interneurons upstream of motor neurons support posture maintenance^28^. Because APN2 rests at a high membrane potential, we propose that its reduced excitatory influence during flight (Figure 1E) contributes to this flight-specific antennal posture by reducing excitation of downstream antennal motor neurons. Consistent with this idea, optogenetic inaction of APN2 produced a subtle but significant dorsal shift in antennal posture (Figure 4E), supporting a role for APN2 hyperpolarization in regulating antennal posture during flight.

### Dynamic integration of antennal mechanosensory signals by APN2

Despite its tonic decrease in activity during flight (Figure 1E), APN2 continued to respond to external, airflow-mediated mechanical displacements of the antennae with graded hyperpolarization (Figure 2B). The magnitude, timing, and rate of APN2 responses to airflow were comparable during quiescence and flight when the JONs were intact (Figures 2F-H), demonstrating that APN2 preserves stimulus-evoked hyperpolarization during quiescence and flight, allowing the animal to remain sensitive to external mechanical cues even during flight-induced tonic inhibition. Consistent with this idea, antennal position distributions during airflow were broader during flight than during quiescence (Figures 2D-E). This increased variability suggests that reduced APN2 activity may permit greater antennal movement while preserving sensory responses, thereby supporting enhanced mechanosensory sampling during active movements.

Although APN2 remained responsive to airflow in the absence of JON input (Figure 2C), previous work showed that its wind responses are abolished when the ipsilateral antenna is removed^31^, suggesting that additional antennal mechanosensors besides JONs contribute to APN2 activity. Using connectomics data, we show that APN2 lies downstream of antennal bristle sensory neurons (Figure 5Aii). The 2-4 synaptic hops between bristle sensory neurons and APN2 (Figure 5Aii) could explain the delayed response timing between the onset of airflow compared to the onset of APN2 response when JONs are blocked (Figure 2E). The pathways from antennal bristles to APN2 exhibit a roughly equal proportion of excitatory and inhibitory connections (Figure 5Biii), providing a rationale for the bimodal response to airflow we observed in many APN2 cells when the JONs were blocked (Figure 2C). This balance also emphasizes the diversity of APN2 responses, where individual neurons may receive different combinations of sensory inputs (Figure 5A). By encoding sensory signals from both JONs and antennal bristles, and with its established role in encoding wind direction^31^, APN2 is well positioned to integrate mechanosensory cues from the antennae to support mechanosensory-guided behaviors, like steering or groundspeed regulation.

### APN2 encodes active antennal movements

Even in the absence of external airflow, APN2 responded when the antennae moved spontaneously (Figure 3A), demonstrating sensitivity to self-generated motor signals and a role beyond detecting external mechanical cues^31^. Not only did APN2 respond to active movements of the ipsilateral antenna, but it also encoded movements of the contralateral antenna (Figure 3A), indicating a role in integrating motor-related signals from both antennae. The lack of delay in APN2’s response between ipsilateral and contralateral antennal movements suggests that it lies downstream of higher-order motor circuits that lie upstream of both APN2 and antennal motor neurons. Connectomic analyses revealed neurons projecting to both APN2 and antennal premotor neurons on ipsilateral and contralateral sides (Figure 5D), providing candidates for investigating how antennal motor commands influence APN2 and its mechanosensory signaling (Table S2).

When JON input was removed, APN2 still exhibited graded changes in membrane potential during active antennal movements (Figure 3B). Although this difference was not statistically significant, APN2 was, on average, more responsive during antennal movement than stillness (Figure 3C), suggesting a role for APN2 in antennal motor control. Even modest changes in membrane potential – as small as 2 mV in nonspiking interneurons – are sufficient to drive neurotransmitter release onto downstream motor neurons^25,41^, supporting the plausibility of APN2’s influence on the antennal motor system. Further, the increase in antennal movements when mechanosensory input was blocked (Figure 3B) indicates that antennal activity during quiescence is shaped by feedback from mechanosensory signals, highlighting APN2’s potential role in integrating this feedback to regulate active movements. Our confidence in this interpretation is strengthened by our optogenetic results that show that activating APN2 elicits antennal movements (Figures 4F-G). The connectome supports this functional role, as we identified pathways as short as two synapses between APN2 and antennal motor neurons (Figure 5Ai).

Beyond response timing and circuit position, the polarity of APN2 responses provides additional insight into its sensory-motor function. Notably, APN2 exhibited tonic hyperpolarizing or depolarizing responses to similar active antennal movements, both in the presence and absence of JON input (Figure 3A-B). We believe this variation reflects differences in the presynaptic partners of individual APN2 cells. These differences may be functionally meaningful, as supported by our connectomic analyses showing that APN2 projects to two different antennal motor neurons (Figure 5, Table S2). This bimodal encoding could, for example, help coordinate antagonistic motor outputs. Our interpretation aligns with observations in other arthropods, where individual nonspiking interneurons can excite motor neurons to one muscle while simultaneously inhibiting motor neurons of the antagonist, thereby enabling precise motor control^24,26,41,42,43^. By encoding the excitatory and inhibitory inputs to motor neurons, APN2 is well-positioned to integrate mechanosensory and higher-order motor signals, enabling flexible control of antennal movements during active behavior.

## Summary

Our results provide evidence that APN2 contributes to antennal sensory-motor integration. By balancing input from multiple mechanosensors with higher-order motor signals, APN2 integrates sensory and motor-related commands to direct motor output. These findings provide a framework for exploring other nonspiking interneurons for sensory-motor integration. Through connectomic analyses, we also identify candidate pathways for future studies aimed at understanding how sensory-motor circuits guide behavior.

## Materials and Methods

### Fly husbandry

We used adult female *Drosophila melanogaster,* between the ages of 2 and 10 days old, for all experiments. All flies were maintained in an incubator at 25°C on a 12-hour light/dark cycle and raised on standard cornmeal molasses food. For both electrophysiology and behavior experiments we used a GAL4 driver line that targets all 9 APN2 cells in each hemisphere (Figure S4) and has been used in two previous studies^31,32^. To visualize cells for electrophysiology recordings, we expressed cytoplasmic GFP (*UAS-10xGFP*) under the control of our APN2-GAL4 driver (*R24C06-GAL4*). For optogenetic behavior experiments, we used flies expressing either *UAS-GtACR1* or *UAS-CsChrimson* under the control of our APN2-GAL4 driver (*R24C06-GAL4/UAS-GtACR1 or R24C06-GAL4/UAS-CsChrimson,* respectively). Control flies were *Canton-S x UAS-GtACR1 and Canton-S x UAS-CsChrimson*. All flies used in optogenetic experiments were placed on hydrated potato flakes (Bob’s Red Mill) supplemented with 35 mM all-trans retinal (Sigma) 24-72 hours prior to experimentation.

### Immunohistochemistry and anatomy

To validate the number of APN2 cells targeted by our APN2 driver line (*24C06-GAL4*), we dissected the fly’s brain in PBS and used immunohistochemistry to amplify the expression of the mVenus fluorophores in *24C06-GAL4>UAS-CsChrimson* flies. First, we fixed the brain for 15 min in a PBS solution containing 4% paraformaldehyde. We then washed the brain three times before incubating it in blocking solution (5% normal goat serum in PBST) for 20 minutes. Next, we incubated the brain in primary antibody solution containing 1:10 mouse anti-bruchpilot (nc82, Developmental Studies Hybridoma Bank) and 1:1000 rabbit anti-GFP (Invitrogen #A6455) in block. After 24 hours at room temperature, we washed the brain three times in PBST. Then, we incubated the brain in secondary antibody solution containing 1:250 anti-mouse Alexa Fluor 633 (Invitrogen #A21052) and 1:250 anti-rabbit Alexa Fluor 488 (Invitrogen #A11034) in block, again for 24 hours at room temperature. We washed the brain three times in PBST before mounting it in vectashield (Vector Labs H-1000). The slide was stored at 4°C until imaging.

We imaged the brain at 20x magnification on a Zeiss LSM 880 confocal microscope with a 20x objective (Plan-Apochromat 20x/0.80 M27 0.55mm). All brains were imaged at 1.04 µM depth resolution. The final images presented in our figures are maximum z-projections.

### Electrophysiology

Using previously described methods and solutions, we performed whole cell patch clamp recordings in tethered flies^45,46^. We briefly cold-anesthetized flies and removed the middle and hind legs at the femur-trochanter joint and the front legs at the base of the tibia to prevent interference with antenna tracking. We then used UV-curing glue (Wayin UV Resin Hard) to attach the fly to a plastic holder. The holders were 3D printed (Vanderbilt University Wondry) and attached to folded metal cutouts (Etchit) with UV-curing glue, as previously described^47^.

To prevent excess brain movement, we immobilized the proboscis with UV glue and secured it to the prothoracic femurs. Then, under saline, we used a small hypodermic needle (30G) to remove the cuticle spanning from the ocelli to just dorsal to the antennae. Using fine forceps, we gently removed the trachea on the surface of the brain just over the region where our cell bodies resided. We broke through the sheath of the brain and cleaned the area around the cell bodies of interest using a small amount of collagenase (5% in extracellular saline; Worthington Biochemical Corporation Collagenase Type 4), which we applied with a fine-tip electrode (approximately 5-10mm in diameter; item #TW150F-3, World Precision Instruments) at room temperature. During all experiments, we perfused 275 mOsm extracellular saline composed of 103 mM NaCl, 3 mM KCl, 5 mM TES, 10 mM trehalose dihydrate, 10 mM glucose, 2 mM sucrose, 26 mM NaHCO_3_, 1 mM NaH_2_PO_4_, 1.5 mM CaCl_2_2H_2_O, and 4 mM MgCl_2_6H_2_O over the brain. We bubbled the saline with 95% O_2_/ 5% CO_2_ to get the pH near 7.3.

We used 6-11 MΩ thick-walled glass pipettes (Item #1B150F-3, World Precision Instruments) for whole cell patch clamp recordings, which we pulled using a Sutter P-1000 puller. Our intracellular saline contained 140 mM K-aspartate, 1mM KCl, 10 mM HEPES, 1 mM EGTA, 0.5 mM Na_3_GTP, 4 mM MgATP. We also added 0.5% biocytin hydrazide to visualize neural processes post-hoc (data not shown). Using *R24C06-GAL4/UAS-10xGFP*, we pseudorandomly targeted one of the 9 cell bodies in the left hemisphere for all recordings.

We used an A-M Systems Model 2400 patch clamp amplifier equipped with a 10 G Ω /100 MΩ headstage set to low gain (100 mV/nA) in our electrophysiology experiments. APN2 neurons had an average membrane potential of −34 mV, which includes a measured liquid junction potential estimate of −13 mV^45^.

All recordings were performed in an arena enclosed by a light and wind-proof faraday cage, which we achieved by attaching foam-core boards to the outside of the cage and taping the edges (using black gaffer’s tape) to prevent external airflow. We used two Basler Ace cameras (Edmund Optics), each with a 44mm/3.00x lens (Infinistix) and a 2x doubler tube lens (Edmund Optics), with a sample rate of 100 Hz and a resolution of 640×480 pixels for lateral tracking of both antennae (see Antennal Tracking). The arena also included a manual air puffer (see Experimental Design) and a wingbeat detector (see Flight Detection). All stimuli were sent and controlled using custom MATLAB scripts combined with a data acquisition card and BNC breakout board (PCIE-6323 and BNC-2090A, National Instruments). All data, besides the video data, were acquired at 10 kHz. When we made comparisons with antennal data, we down sampled the data to 100 Hz.

### Optogenetic behavior

Before pin-tethering flies for behavior experiments, we briefly cold-anesthetized them and severed each leg at the femur-trochanter joint. We used an Allied Vision Guppy Pro (Edmund Optics) camera with a 44mm/3.00x lens (Infinistix) running at 60Hz with a resolution of 640×480 pixels for tracking the fly’s right antennae from a lateral view (see Antennal Tracking). Additionally, we installed a long-pass infrared filter (Edmund Optics) on the camera lens to eliminate lighting artifacts from the optogenetic light stimulus in the recorded video. We simultaneously recorded wingbeats using a microphone positioned perpendicular to the fly’s left wing (see Flight Detection). The arena also included fiber optics with 505 nm or 590 nm LEDs that we aimed at the fly’s head capsule for optogenetic inactivation or activation, respectively (Thorlabs). All optogenetic behavior experiments were conducted in a light-proof and sound-reduced arena, which we achieved by enclosing the arena with foam-core boards layered with 1 inch sound-proofing foam (Hemrly) and covering it with a bead-filled weighted blanket (15 lbs; TreeCube). Stimuli were sent and controlled using custom MATLAB scripts combined with a data acquisition card and BNC breakout board (PCIE-6363 and BNC-2110, National Instruments). All data was acquired and analyzed at 60 Hz.

We presented each fly with 50 experimental trials where 40 trials consisted of a 4 second light stimulus buffered by a 2 s pre-stimulus and 4 s post-stimulus window, and 10 random trials in which there was no light stimulus. The length and intensity of the optogenetic light stimulus (inactivation: 7.75 mW/cm^2^, activation: 2.6 mW/cm^2^) follows previous work demonstrating little adaptation in these cells despite a prolonged stimulus period^31^. For each stimulus-containing trial, we compared the average antennal position during the last 2 s of the stimulus window to the 2 s pre-stimulus window. All average antennal responses to the optogenetic light stimulus in Figure 4 are presented as averages across individual flies. For inactivation experiments, we collected data from 10 control flies and 11 experimental flies, whereas for our activation experiments, we collected data from 15 control flies and 15 experimental flies. We only included trials where the fly was either flying or quiescent for the entire trial. Trials where the fly either started or stopped flying during the 10 s trial were excluded from analysis.

### Antennal tracking

We used fiber optics equipped with infrared LEDs (850 nm; Thorlabs) to illuminate the fly during video acquisition, and DeepLabCut^48^ (version 2.3.6) to track recorded antennal movements. For tracking with DeepLabCut, we used a ResNet-50 neural network, trained over 500,000 iterations with a training-to-test fraction of 0.8 in all experiments.

For electrophysiology experiments we trained a DeepLabCut network for each camera view and each antennal condition (Table 1).

**Table 1.**
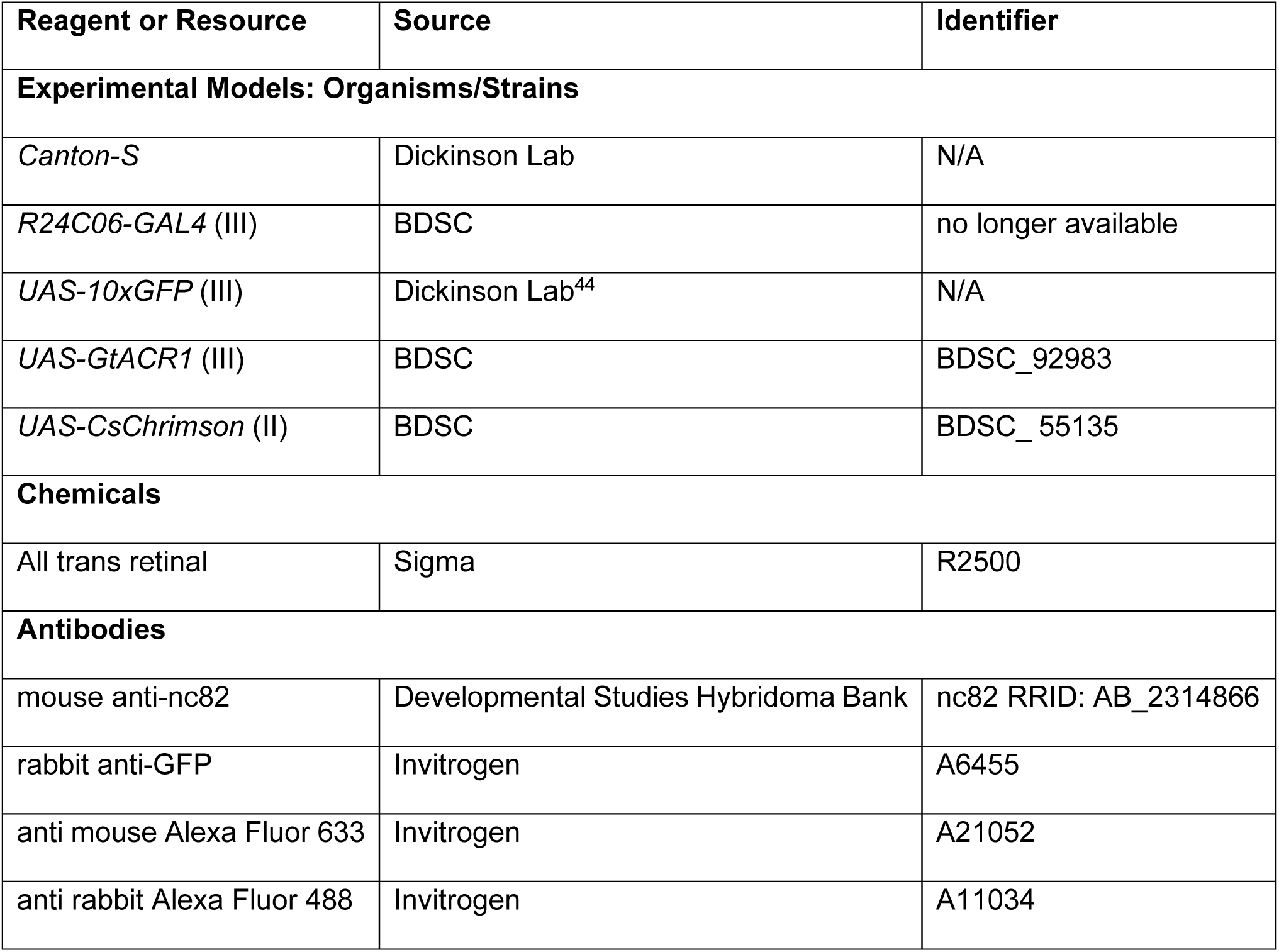
Reagents and resources used in this study. We used transgenic fly lines in our electrophysiology and behavior experiments. For our optogenetic experiments, we used all trans retinal.

| Reagent or Resource | Source | Identifier |
| --- | --- | --- |

| Experimental Models: Organisms/Strains |  |  |
| --- | --- | --- |
| *Canton-S* | Dickinson Lab | N/A |
| *R24C06-GAL4* (III) | BDSC | no longer available |
| *UAS-10xGFP* (III) | Dickinson Lab<sup>44</sup> | N/A |
| *UAS-GtACR1* (III) | BDSC | BDSC_92983 |
| *UAS-CsChrimson* (II) | BDSC | BDSC_55135 |
| Chemicals |  |  |
| All trans retinal | Sigma | R2500 |
| Antibodies |  |  |
| mouse anti-nc82 | Developmental Studies Hybridoma Bank | nc82 RRID: AB_2314866 |
| rabbit anti-GFP | Invitrogen | A6455 |
| anti mouse Alexa Fluor 633 | Invitrogen | A21052 |
| anti rabbit Alexa Fluor 488 | Invitrogen | A11034 |

**Table 2.**
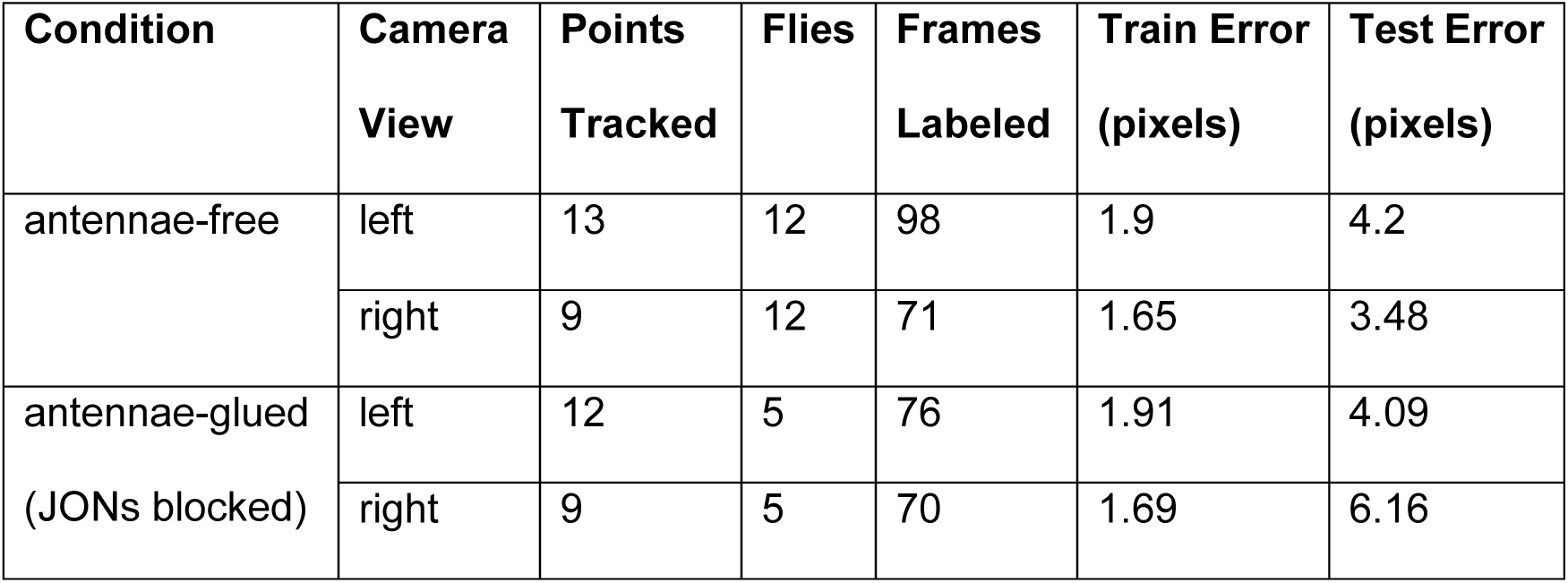
DeepLabCut Network Properties. We trained 4 networks, 1 for each antennal view (ipsilateral and contralateral) within each antennal condition (free and glued).

To quantify the position of the antenna during each trial we compared the antennal axis (the angle of the top-most attachment site of the first antennal segment to the head capsule down to the tip of the third segment) to the vertical axis of the video frame. For the data in Figure 2, we averaged all trials from the ipsilateral antenna that was collected from each APN2 cell to obtain the average airflow induced antennal deflection over time in each animal. We then averaged all animals together to get the average antennal response within each condition. Data in Figure 3, however, displays all spontaneous movements for the ipsilateral and contralateral antenna.

For optogenetic experiments, we trained a DeepLabCut network using 100 frames from 24 flies across genotypes and experimental designs. We tracked 14 points across the head and antennae of the fly. The trained network had a training error of 1.66 pixels and a testing error of 3.92 pixels. For every fly, we defined a reference axis using stationary points at the base of hairs above and below the eye consistently across animals. To quantify antennal position during each trial, we compared the angle between the base of the arista and the tip of the third antennal segment to the reference axis. We averaged all trials collected from each fly to obtain the average antennal deflection over time in each animal and then averaged all the animals together to get the average antennal response.

### Flight detection

To detect wing movement in our electrophysiology experiments, we used a liquid light guide coupled to a custom circuit board to continuously collect infrared light reflected off the wings, as previously described^49^. We extracted the amplitude of the wingbeat envelope using a Hilbert transform. To reduce high-frequency noise, the extracted envelope was smoothed using a low-pass Butterworth filter (6 Hz cutoff, 10 kHz sampling rate). Flight bouts were then identified by applying a custom threshold to the smoothed signal. The resulting wingbeat signal was categorized as a flight bout if its duration was 150 ms or longer.

We compared the neural responses of APN2 during bouts of quiescence and/or bouts of flight. All reported flight bouts in Figures 1-3 were initiated with a brief pulse of air, so we excluded the 500 ms period of data following each air puff occurrence from our analyses. We also excluded the 100 ms period of data following every bout of flight to avoid any state transition effects and ensure stable classifications of flight and quiescence. With the resulting data, we then identified bouts of quiescence and flight that either excluded (Figures 1 and 3) or included (Figure 2) air puffs.

In our optogenetic behavioral experiments in Figure 4, we used a miniature lavalier microphone to monitor wing activity (Countryman Audio, B6P4FF05B), similar to previously described methods^50^. We detected wingbeats by low-pass exponential smoothing the signal (α = 0.9995) followed by finite impulse response filtering to remove residual high-frequency noise. This allowed us to extract a smoothed wingbeat envelope while filtering out individual wingbeat oscillations. Flight bouts were determined by applying a fixed amplitude threshold to the resulting envelope. Trials where the fly only flew for a portion of the experiment were not included in our analyses, and only fully flying trials or fully quiescent trials were analyzed.

### Blocking JONs

We also investigated the influence of JONs activity on APN2 responses by placing a small drop of glue between the second and third antennal segments to block JON sensation^33^. In all glued experiments (Figures 1-3), we glued both antennae, and we left both antennae unmanipulated in all antennae-free experiments.

In Figure 1, we averaged the membrane potential of APN2 across all bouts of quiescence or flight, both when the antennae were free and when they were glued. All average recordings in Figure 1 were obtained from 21 individual APN2 cells across 21 flies. The presented averages were calculated from stable recordings that contained a range of 2-124 bouts and an average of 39 bouts.

### Airflow stimulus

To investigate sensory responses to airflow during our electrophysiological experiments (Figure 2), we used a manual air puffer – a 10 mL plastic syringe connected to output tubing (3.18 mm ID/6.35 mm OD). We positioned the puffer 35 mm from the fly at a 45 deg angle, contralateral to the side of the brain we recorded from, to deliver mechanical stimuli to the fly. The air puffer was connected to an airflow sensor (Honeywell Sensing Solutions AWM3300V), whose signal we recorded at a sampling rate of 10kHz. Air puff occurrences were detected using peak-based thresholding (minimum peak height = 2 V, prominence = 1 V, width = 30 ms) and further filtered to focus on the air puffs that were less than 400 ms in length. We included an additional 100 ms of data beyond the threshold-defined end of each air puff to capture associated antennal and neural responses. We used a hot wire anemometer (MiniCTA with 55P11 probe, Dantec Dynamics), positioned where the fly would be positioned during recordings, to measure the timing of stimulus onset (gold, dashed line; Figure 2A-C). We measured no statistically significant difference between individual manual air puffs (Figure S1).

Occasionally, the air puffs initiated flight, producing two responses: sustained flight that persisted after the puff, or flight that ceased once the airflow ended. APN2’s response to these two subcategories of flight did not differ significantly (data not shown; Wilcoxon Signed-Rank test; antenna free comparison: p = 0.812, antennae glued comparison: p = 0.250), so the flight data we present in Figure 2 encompasses both subgroups. All average recordings in Figure 2 were obtained from 29 individual APN2 cells across 26 different flies. The presented averages were calculated from stable recordings that contained a range of 1-82 puffs and an average of 25 puffs per recorded cell.

### Active antennal movements

To identify spontaneous active antennal movements in our videos tracked with DeepLabCut, we first applied a second-order Butterworth low-pass filter with a 1Hz cutoff frequency to the ipsilateral and contralateral antennal traces. We then identified prominent peaks (scipy.signal.find_peaks; prominence of 3 deg) of the absolute value of each filtered antennal trace during quiescence; we performed this detection both when the antennae were free and when they were glued, to detect trials with candidate movements. We then labeled all antennal movements that were more than 10 degrees in magnitude and ranged from 80 to 110 ms in duration by manually marking the onset and offset of movement. We extracted each start and stop index with an additional 20 ms prior to the onset and following the offset of each spontaneous bout. These same indices were used to extract the corresponding APN2 activity for each bout of movement. All APN2 recordings in Figure 3 are presented as individual responses across 18 APN2 cells for each antenna – all of which were baseline subtracted relative to the first datapoint in the trace.

### Connectomics

All connectomic analyses were performed using a combination of custom Python and MATLAB scripts. We used version 783 of the FAFB dataset^34,35^, which was downloaded from: https://codex.flywire.ai/api/download?dataset=fafb.

Cell type definitions of JONs (‘JO’), antennal bristles (‘BM_ant’) and all other cell types not explicitly mentioned below are based on version 783 annotations^34,35,37,51,52^. In total, seven APN2 neurons were identified in each hemisphere; two neurons remain without a mirror pair. Using the classifications and connections datasets, we indexed out all 14 identified APN2 neurons (Table 3) with their inputs, and all the associated metadata.

**Table 3.** Identified APN2 neurons in the FAFB connectome. We identified 14 APN2 cells in the connectome; 12 of which are bilaterally symmetric, and 2 remaining without an identified mirror pair.

| Cell ID | Cell Name | Side | Mirror Pair |
| --- | --- | --- | --- |
| 720575940614190242 | SAD.182 | Left | A |
| 720575940635960292 | WED.102 | Right |  |
| 720575940629552170 | SAD.50 | Left | B |
| 720575940627027087 | AMMC.WED.11 | Right |  |
| 720575940606919602 | SAD.IPS.57 | Left | C |
| 720575940625952002 | IPS.418 | Right |  |
| 720575940629394232 | SAD.IPS.30 | Left | D |
| 720575940613641794 | AMMC.IPS.2 | Right |  |
| 720575940650920569 | SAD.64 | Left | E |
| 720575940618772719 | AMMC.WED.17 | Right |  |
| 720575940603930046 | AMMC.SAD.5 | Left | F |
| 720575940620833974 | AMMC.WED.12 | Right |  |
| 720575940630214564 | SAD.51 | Left | N/A |
| 720575940625597317 | AMMC.WED.19 | Right |  |

We analyzed pathways from the JONs, antennal bristles, and any additional inhibitory cells to APN2, and from APN2 to antennal motor neurons. Pathways were identified using a depth-first search algorithm to find all possible connections up to four hops (three interneurons) between defined start and stop nodes. For pathways between the JONs and APN2, we restricted our search to 2 hops (1 interneuron). Synaptic connections between an upstream neuron onto a given downstream partner with fewer than five synapses were excluded from analyses, following the Codex synaptic partner criteria^34,37^. The resulting CSV was analyzed with custom MATLAB scripts. We sorted identified nodes based on which side the cell resides on in the connectome (left or right), and if its predicted neurotransmitter type is excitatory (ACH) or inhibitory (GABA or glutamate). Some cells were classified as central, instead of left or right, and they are indicated in the pathways in which they occurred. To identify candidate cells belonging to the ‘higher order motor’ node, we searched the connectome for matching inputs upstream of APN2 and the cells we identified as premotor neurons. In this search, we excluded recurrent APN2 connections and connections with the JONs. Each directed graph was plotted with user-set parameters to yield a weighted network graph containing neuron count, synapse counts, and synaptic weights. Unless otherwise indicated, synaptic inputs and outputs from the left and right hemispheres were pooled for analysis; hemispheres were analyzed separately only in plots where left and right are explicitly distinguished.

### Experimental design and statistical analysis

All data was collected in MATLAB 2024b and analyzed using custom python and MATLAB scripts. Alpha values were set at 0.05 for statistical tests, with * representing <0.05, ** representing ≤0.01, and *** representing ≤0.001. Across all experiments, we used Shapiro–Wilk’s test to test the normality of data (python: scipy.stats.shapiro; MATLAB: swtest^53^). We also used Levene’s test to test for equal variance (scipy.stats.levene).

Our electrophysiology experiments compared two categorical variables: flight state (quiescent versus flight) and antennal condition (free versus glued). All flight state comparisons were paired because we only analyzed data from flies that flew. When the data was normally distributed, we used paired t-tests (Python: scipy.stats.ttest_rel; MATLAB: ttest) to compare flight state; for non-normal data, we used the Wilcoxon Signed-Rank test (Python: scipy.stats.wilcoxon; MATLAB: signrank). All comparisons made across antennal conditions were independent. When the data was normally distributed, we used independent t-tests (Python: scipy.stats.ttest_ind; MATLAB: ttest2) to compare antennal conditions, but when the data was not normally distributed, we used the Mann-Whitney U test (Python: scipy.stats.mannwhitneyu; MATLAB: ranksum).

Two categorical variables were also compared in our optogenetic behavior experiments: flight state (quiescent versus flight) and experimental condition (control versus experimental). All comparisons were unpaired, because some flies did not fly, whereas other flies flew for the duration of the experiment. When the data was normally distributed, we used independent t-tests to compare the effect of the optogenetic stimulus versus the control, but when the data was not normally distributed, we used the Mann Whitney U test (Figure 4).

## Code accessibility

The authors would like to thank Seaborn, SciPy, NumPy, Pandas, Matplotlib, and emd for their contributions to free and open-sourced software. All the data and code generated in this study is available on GitHub (https://github.com/suverlab/NunnEtAl2026).

## Acknowledgements

This work was funded in part by grants awarded to M.P.S. from the National Institutes of Health Brain Initiative (R00NS114179, R01NS140174, and U01NS131438) and to O.M.N. from the National Institute of Neurological Disorders and Stroke (F31NS141622). Stocks obtained from the Bloomington Drosophila Stock Center (NIH P40OD018537) were used in this study. Confocal experiments were performed in part with the Vanderbilt Cell Imaging Shared Resource (supported by NIH grants S10OD021630, CA68485, DK20593, DK58404, DK59637, and EY08126). The authors would like to thank Kelly Collins at Trew Audio and Dr. Hokto Kazama for advice regarding our microphone set up for detecting wingbeats, and Dr. Alexandre Tiriac for helpful discussion.

## Supporting Information

**Figure S1.**
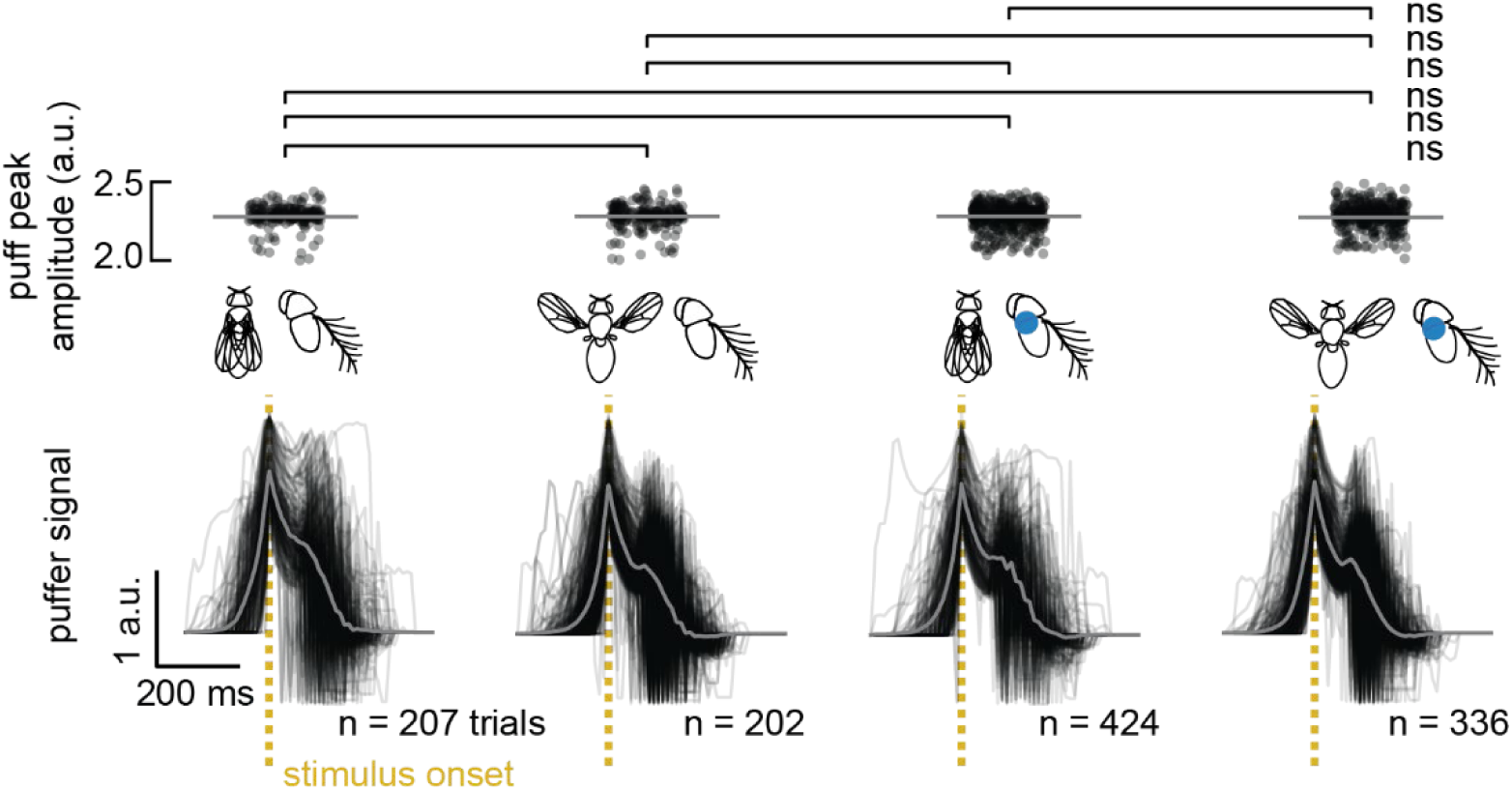
The peak amplitudes of manually delivered air puffs were consistent across conditions. (Bottom) The air puff detected in each trial in Figure 2 is plotted in black, respective of the condition it occurred in, and with the average trace plotted in gray. All traces were aligned by their highest peak, which is representative of the time point in which the air puff reached the fly (‘stimulus onset’, gold dashed line). (Top) Each trial’s puff peak amplitude is plotted as a black dot, with the gray bar representing the conditional average across all air puffs. No significant difference in the amplitude of air puff peaks was observed between conditions (Mann-Whitney U; all p>0.05).

**Figure S2.**
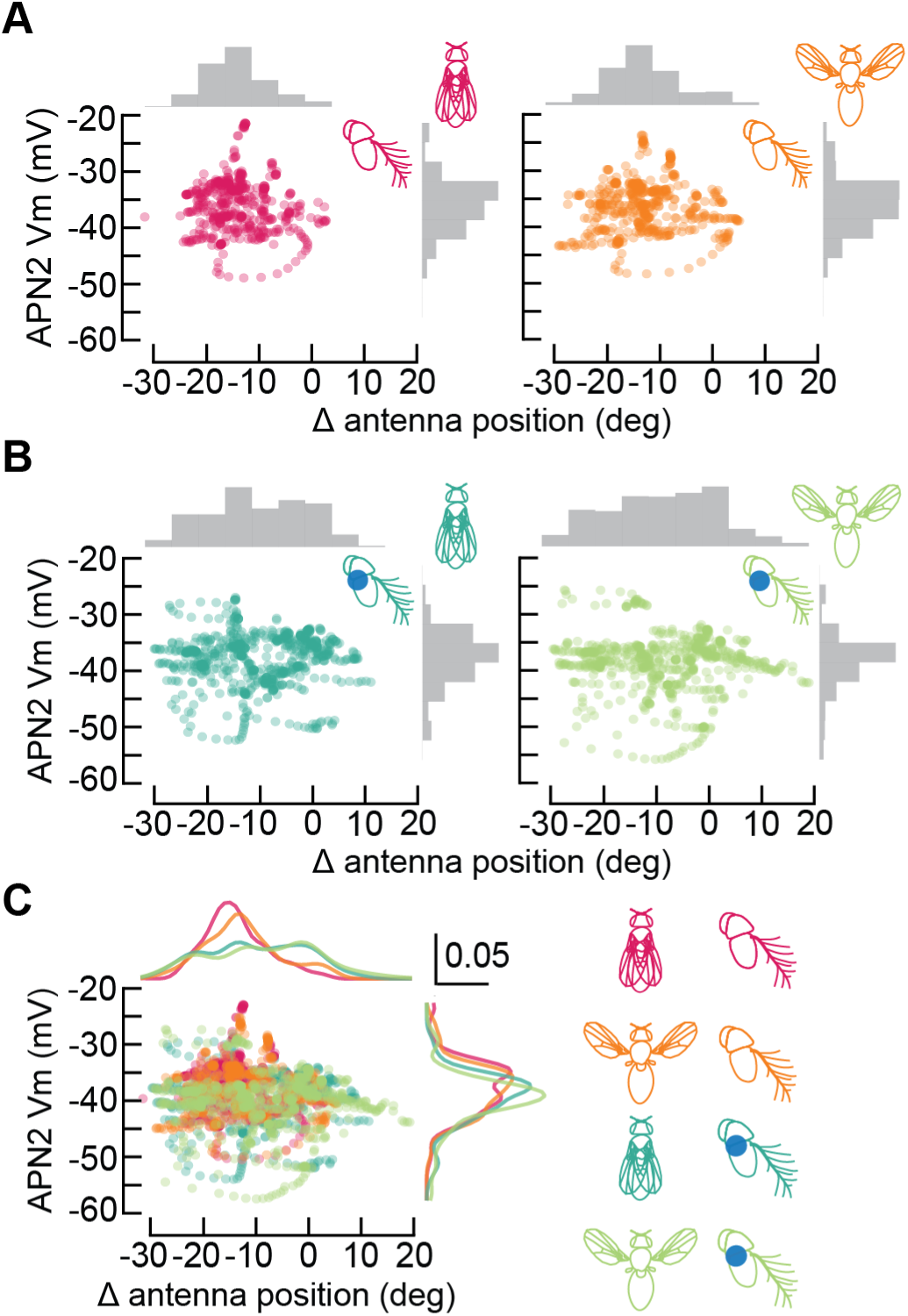
Antennal movements are higher in distribution in response to air puffs when JONs are blocked. A) Relationship between antennal position and APN2 membrane potential during air puffs with free antennae in quiescent (left) and flying (right) flies. Each dot represents the average air puff response of an individual fly (data shown in Figure 2B-C), with colors denoting experimental conditions. Distributions of antennal position and APN2 membrane potential are shown as histograms along their respective axes. The distribution of antennal position increases in variance in flying vs. nonflying flies (pairwise Levene’s test; p < 0.001) B) Similar to (A), but when the JONs are mechanically blocked. Blocking JONs increased the variability of antennal position during air puffs in both quiescent and flying flies (p < 0.001). The distribution of antennal position remained different in flying vs. nonflying flies, even when JONs were blocked (p =0.012). C) The distributions of antennal position and APN2 membrane potential are shown as density curves along the respective axes. The correlation between APN2 membrane potential and the ipsilateral antenna position shows no clear trend. The distribution of antennal position has significant variability across all conditions (Levene’s test; p < 0.001), but the distribution of APN2 membrane potential remains relatively consistent (p = 0.144).

**Figure S3.**
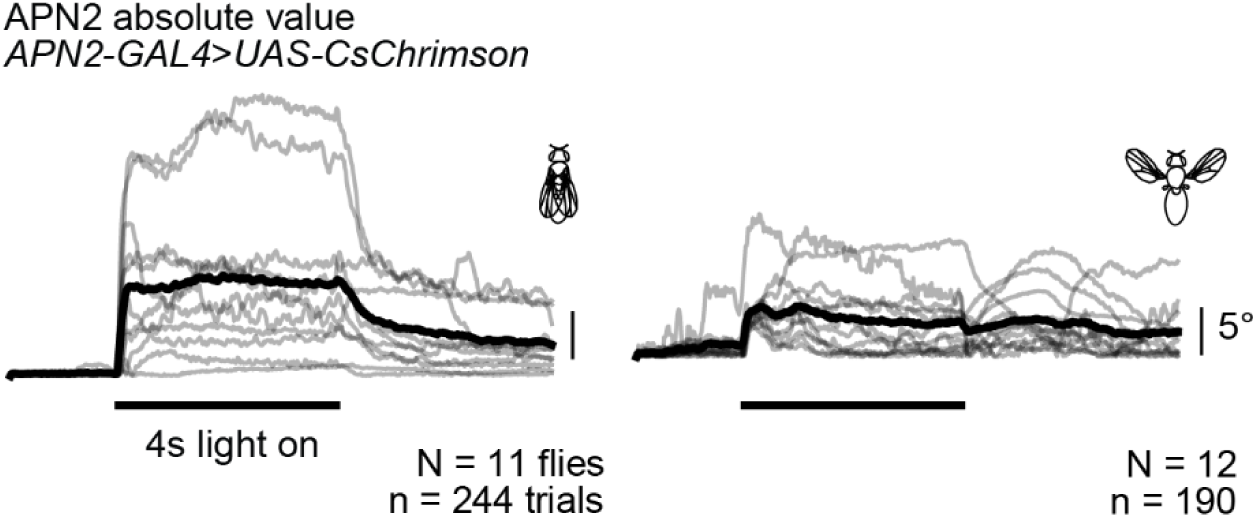
Activation of APN2 drives antennal movements. Comparison of the absolute value of the average antennal response to optogenetic activation of APN2 in quiescent (left) and flying (right) flies. Each gray trace is the average of all trials for each fly, and the black traces represent the average response across all flies.

**Figure S4.**
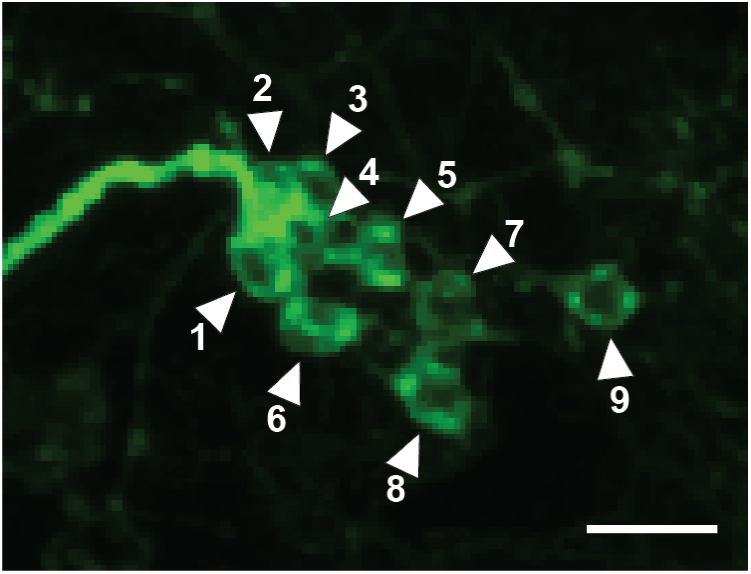
Nine APN2 cells are labeled by the genetic driver line *24C06-GAL4*. Confocal maximum projection image of *24C06-GAL4>CsChrimson*, with the mVenus tagged *UAS-CsChrimson* expression in green. White arrowheads point to APN2 cell bodies and are numerically numbered from medial to lateral. Scale bar is 10µm.

---

**Table S1.** Number of neurons identified in each node of the sensory-motor circuit involving APN2. The “Node” column uses nomenclature from Figure 5 but separates the nodes further into excitatory and inhibitory groups. The number of neurons belonging to each node are also separated into left and right, depending on which side of the brain they are found on. Over 3,225 neurons were identified in this network.

| Node | Number of Neurons |  |
| --- | --- | --- |
|  | Left | Right |
| JONs | 570 | 473 |
| JONs sensory interneurons (excitatory) | 330 | 251 |
| JONs sensory interneurons (inhibitory) | 284 | 259 |
| JONs central sensory interneurons (inhibitory) | 16 |  |
| antennal bristles | 61 | 61 |
| antennal bristle sensory interneurons (excitatory) | 107 | 80 |
| antennal bristle sensory interneurons (inhibitory) | 84 | 79 |
| APN2 | 7 | 7 |
| flight-specific inhibitory cells | 3 | 1 |
| premotor interneurons (excitatory) | 88 | 83 |
| premotor interneurons (inhibitory) | 96 | 91 |
| AMNs | 3 | 1 |
| higher-order motor (excitatory) | 13 | 9 |
| higher-order motor (inhibitory) | 63 | 62 |

**Table S2.** Cell types for future investigation into antennal sensory-motor integration. We provide the cell IDs and names of a subset of neurons identified in the sensory-motor circuit that APN2 contributes to. The “Node” column uses nomenclature from Figure 5. All other information was obtained from our connectomics analyses.

| Node | ID | Name | Side | NT type<br>(predicted) |
| --- | --- | --- | --- | --- |
| AMNs | 720575940621351412 | GNG.638 | Left | N/A |
|  | 720575940622235930 | GNG.842 | Right | N/A |
|  | 720575940629183643 | GNG.647 | Left | N/A |
|  | 720575940639021757 | GNG.1350 | Left | N/A |
| flight-specific<br>inhibition | 720575940610640482 | SPS.1 | Right | N/A |
|  | 720575940625484069 | GNG.AMMC.17 | Left | GABA |
|  | 720575940626359368 | IPS.AMMC.16 | Left | GABA |
|  | 720575940629486714 | SPS.IPS.1 | Left | GABA |
| higher-order<br>motor | 720575940613869777 | AL.197 | Left | ACH |
|  | 720575940630907434 | SAD.21 | Left | ACH |
|  | 720575940615393959 | IPS.51 | Left | GLUT |
|  | 720575940633803676 | IPS.57 | Right | GLUT |
|  | 720575940628785468 | SAD.27 | Right | GLUT |

